# Common and distinct roles of AMPKγ isoforms in small-molecule activator-stimulated glucose uptake in mouse skeletal muscle

**DOI:** 10.1101/2025.09.26.678768

**Authors:** Dipsikha Biswas, Ever Espino-Gonzalez, Danial Ahwazi, Jordana B. Freemantle, Charline Jomard, Jonas Brorson, Agnete N. Schou, Jean Farup, Julien Gondin, Jesper Just, Marc Foretz, Jonas T. Treebak, Marianne Agerholm, Kei Sakamoto

## Abstract

**Objective:** Small-molecule activators targeting the allosteric drug and metabolite (ADaM) site of AMPK enhance insulin-independent glucose uptake in skeletal muscle and lower glucose in preclinical models of hyperglycemia. The regulatory AMPKγ subunit plays a central role in energy sensing. While the skeletal muscle-selective γ3 isoform is essential for AMP/ZMP-induced glucose uptake, it is dispensable for ADaM site-binding activators. We hypothesized that the predominant γ1 isoform is required for ADaM site activator-stimulated glucose uptake in skeletal muscle.

**Methods:** Single-nucleus RNA sequencing (snRNA-seq) was performed on mouse and human skeletal muscle mapping AMPK subunit isoform distribution across resident cell types. To determine γ isoform-specific requirements for activator-stimulated glucose uptake, skeletal muscle-specific inducible AMPKγ1/γ3 double knockout (imγ1^-/-^/γ3^-/-^) and single knockout (imγ1⁻^/^⁻ and imγ3⁻^/^⁻) mice were generated*. Ex vivo* glucose uptake was measured following treatment with AICAR (AMP-mimetic) or MK-8722 (ADaM site activator), and *in vivo* MK-8722-induced blood glucose lowering was assessed.

**Results:** snRNA-seq revealed distinct AMPK isoform distribution: γ1 was ubiquitously expressed, whereas γ3 was enriched in glycolytic myofibers in both mouse and human skeletal muscle. *Ex vivo*, glucose uptake stimulated by either AICAR or MK-8722 was abolished in imγ1⁻^/^⁻/γ3⁻^/^⁻ muscle, and MK-8722-induced blood glucose lowering was significantly blunted *in vivo*. AICAR but not MK-8722-stimulated muscle glucose uptake was abolished in imγ3⁻^/^⁻, whereas both activators fully retained effects on glucose uptake and glucose lowering in imγ1⁻^/^⁻ mice.

**Conclusions:** While γ1 predominates in stabilizing the AMPKα2β2γ1 complex, it is dispensable for AMPK activator-stimulated glucose uptake in skeletal muscle, whether mediated via the nucleotide-binding or ADaM site.

## 1. INTRODUCTION

AMP-activated protein kinase (AMPK) is a central energy sensor that plays a pivotal role in maintaining cellular homeostasis through coordinating metabolic signaling pathways in response to acute energetic perturbations, such as during physical exercise and muscle contractions, hypoxia, and mitochondrial poisoning [1; 2]. AMPK functions as a heterotrimeric complex composed of a catalytic α-subunit and regulatory β- and γ-subunits. In both humans and rodents, each subunit has multiple isoforms (α1, -α2; -β1, -β2; -γ1, -γ2, -γ3) encoded by distinct genes (*PRKAA1, PRKAA2; PRKAB1, PRKAB2; PRKAG1, PRKAG2, PRKAG3*), allowing for twelve possible heterotrimeric combinations with tissue- and cell type-specific expression profiles [3; 4]. Compositional analyses of skeletal muscle from humans and mice have shown that AMPKα2β2γ1 and AMPKα2β2γ3 are the predominant complexes, accounting for approximately 85–90% of total AMPK trimers in human vastus lateralis [5] and mouse extensor digitorum longus (EDL) muscle [6], with the α2β2γ3 isoform contributing approximately 20% of this pool. AMPKγ3 is expressed almost exclusively in glycolytic skeletal muscle [7], with negligible expression detected in other tissues, such as in brown adipose tissue [8; 9]. In contrast, AMPKγ1 is ubiquitously expressed across tissues [10]. Although AMPKγ2 expression in skeletal muscle has been reported, evidence supporting the formation of functional γ2-containing AMPK heterotrimers remains limited [5; 6; 11]. However, these estimates are derived from immunoprecipitation and immunoblotting analyses of whole skeletal muscle lysates, which do not provide cell-type–specific resolution.

AMPK has long held promise as a therapeutic target for metabolic diseases [12; 13]. The prodrug 5-aminoimidazole-4-carboxamide riboside (AICAR) is the most widely used activator that intracellularly is converted to 5-aminoimidazole-4-carboxamide ribonucleotide (ZMP), an AMP-analogue [14]. Similar to AMP, binding of ZMP to the γ-subunit activates AMPK by three distinct mechanisms: (i) increased Thr172 phosphorylation by LKB1; (ii) decreased Thr172 dephosphorylation by protein phosphatases; (iii) allosteric activation [1; 15-17]. In skeletal muscle, AICAR promotes plasma membrane localization of GLUT4 partly through phosphorylation of TBC1D1, leading to enhanced glucose uptake in an AMPK dependent and insulin independent mechanism [18–21]. Although acute and chronic AICAR treatment has been reported to exhibit a multitude of metabolic health benefits in preclinical models [22–25], AICAR is not a suitable therapeutic option for humans due to poor oral bioavailability and pharmacokinetics [26]. Furthermore, AICAR is not selective for AMPK [27].

In recent years, several allosteric AMPK activators have been identified/developed that bind to a pocket termed the allosteric drug and metabolite (ADaM) site [28] located at the interface of the α subunit (kinase domain N-lobe) and β subunit (carbohydrate binding module) [29]. Among these, potent and orally available systemic pan-AMPK activators, MK-8722 and PF-739, have demonstrated promising efficacy in reversing hyperglycemia in pre-clinical proof of concept studies through insulin-independent glucose uptake via activation of AMPK in skeletal muscle [30; 31]. Through genetic studies in mice, we and others have demonstrated that AICAR-stimulated glucose uptake in skeletal muscle is specifically mediated by the muscle-specific AMPKα2β2γ3 complex. Deletion of any one of the three subunits, AMPKα2 [23; 32; 33], AMPKβ2 [34; 35], or AMPKγ3 [8; 36; 37], abolishes this response. However, it has also been demonstrated that while ADaM-site activator-mediated glucose uptake in skeletal muscle is AMPK-dependent (*i.e*., glucose uptake was ablated in muscle-specific AMPKα1/α2 double knockout mice), this effect is not affected in AMPKγ3^-/-^ muscle [8; 36]. Considering that the γ1-containing AMPK complex predominates in skeletal muscle [5; 6], we hypothesized that AMPKγ1 is critical for mediating ADaM-site activator-stimulated glucose uptake in this tissue. To test this, we generated a series of inducible and skeletal muscle-specific AMPKγ isoform knockout mice; γ1/γ3 double knockout (imAMPKγ1^-/-^/γ3^-/-^), single imAMPKγ1^-/-^ and imAMPKγ3^-/-^. We then assessed the effects of AICAR and ADaM site-activator MK-8722 on glucose uptake in skeletal muscle in each model.

## 2. MATERIALS AND METHODS

### 2.1. Materials

AICAR was purchased from Apollo Scientific (OR1170T; Bredbury, United Kingdom) and MedChemExpress (HY-13417; USA). MK-8722 was purchased from MedChemExpress (HY-111363; USA). Protein G Sepharose was purchased from Cytiva. Cation-exchange paper (P81) was obtained from SVI Phosphocellulose (https://www.svi.edu.au/resources/phosphocellulose_paper/). [γ-^33^P]-ATP was purchased from Hartmann Analytics (SCF-301-12, Germany). Deoxy-D-glucose, 2-[1,2-^3^H (N)] (NET328250UC) and mannitol, D-[2-^14^C] (NEC852250UC) were purchased from Revvity. The AMARA peptide was obtained from GL Biochem (Shanghai) Ltd. MRC Protein Phosphorylation and Ubiquitylation Unit Reagents and Services. All other reagents were from Merck Millipore Sigma unless otherwise stated. List of primary and secondary antibodies are in the **Supplementary Table 1 and 2**.

### 2.2. Animal ethics and models

Animal experiments were conducted in accordance with the French and European directive 2010/63/EU of the European Parliament and of the Council of the protection of animals used for scientific purposes. Ethical approval was given by the Danish Animal Experiments Inspectorate (license number #2021-15-0201-00884) and the local ethic committee CEEA-55 and the French ministry of research (APAFIS#22109).

Previously generated homozygous single AMPKγ1 floxed allele carrying mice [10], single AMPKγ3 floxed allele carrying mice [8] or double AMPKγ1/AMPKγ3 floxed allele carrying mice were crossed with animals heterozygous for the inducible human α-skeletal actin promoter-driven MCM Cre (HSA Cre MCM) [38]. After induction, Cre^+/-^ mice were knockouts (KO, *i.e.*, single imγ1^-/-^, single imγ3^-/-^, double imγ1^-/-^/imγ3^-/-^) while the Cre^-/-^ littermates were used as controls. All animals used in the experiments were on the C57BL/6 background. Both KO and control animals at the age of 9-14 weeks were dosed orally with 40 mg/kg/day of tamoxifen (Sigma-Aldrich T5648) suspended in sunflower seed oil (Sigma-Aldrich S5007) for three consecutive days. All animals were housed under standard conditions with controlled temperature (22 ± 1°C), 12-h light-dark cycle and received water and chow diet (Altromin 1310 or equivalent SAFE DS D30) ad libitum. Experiments were performed on 15-20-week-old male mice unless stated otherwise.

### 2.3. Single-nucleus RNA sequencing of mouse skeletal muscle

We used single-nucleus RNA sequencing (snRNA-seq) data obtained from a separate, unpublished study focused on muscle regeneration after injury. Twelve-week-old male C57BL/6NCrl mice (Charles River Laboratories) were randomly assigned to an injury protocol consisting of 30 electrically-evoked lengthening contractions or to a control procedure (*i.e.*, no lengthening contractions). Gastrocnemius muscle samples were collected before and 2, 4, 7, 14, and 28 days post injury, as well as 28 days following the control procedure, for snRNA-seq analysis. For the present study, only the uninjured samples from the control group were included (n = 6).

Nuclei isolation was performed as previously described [39]. In brief, the medial part of the gastrocnemius muscle was dissected, minced and transferred to 1.5 mL nonstick tubes containing 1.5 mL of lysis buffer (Pure lysis buffer [L9286-180 mL; Sigma], 0.1% DTT [43816-10 mL; Sigma] and 1% Triton X-100 [T1565-1.7 mL; Sigma]). Samples were resuspended for 5 min on ice and centrifuged for 5 min at 750 x g and 4°C using rotor swing arms. The supernatant was removed, and the pellet was resuspended in 1 mL nuclei buffer (PBS, 10% BSA (SRE0036-25 mL; Sigma), 1M MgCl₂ [M1028-100 mL; Sigma] and 0.1 % Protector RNase inhibitor 40 U/μL [3335399001; Sigma]). Samples were then filtered through a 40 μm strainer (43-10040-40; pluriSelect) into a 1.5 mL tube and incubated on ice for 10 min. After incubation, samples were centrifuged for 5 min at 750 x g and 4°C. The supernatant was removed, and the pellet was resuspended in 100 µL nuclei buffer. Each sample received 1 µL of individual TotalSeq-A antibodies (BioLegend) from 0.5 μg/μL stocks to enable multiplexing. Samples were incubated on ice for 30 min. Following this, 1 mL nuclei buffer was added, and the samples were centrifuged for 10 min at 1,000 x g and 4°C. The supernatant was discarded, and the pellet was resuspended in 200 µl nuclei buffer for fluorescence-activated cell sorting (FACS) and library preparation. snRNA-seq was performed by the Single-Cell Omics platform at the Novo Nordisk Foundation Center for Basic Metabolic Research, University of Copenhagen, using the 10x Genomics Chromium single-cell 3′ reagent kit. Libraries were sequenced on an Illumina NovaSeq 6000.

### 2.4. snRNA-seq raw data processing, clustering and cell type annotation

Binary base call (BCL) sequence files were demultiplexed into FASTQ files using BCL Convert v4.0.3. Reads were pseudoaligned using Salmon Alevin [40] v1.10.3 with the flags -read-geometry 2[1–15] -bc-geometry 1[1–16] -umi-geometry 1[17–26] set. RNA libraries were pseudoaligned to the GENCODE [41] vM33 reference transcriptome, distinguishing between spliced and unspliced transcripts. Alevin-fry [42] v0.9.0 was used to quantify both the RNA and HTO libraries. Quantitated libraries were processed with Seurat [43] v5.0.1. Empty droplets were detected by the barcodeRanks function from the DropletUtils [44] package. HTOs were normalized using the NormalizeData function with the ‘CLR’ normalization method. A Gaussian mixture model with two components was fitted to each HTO distribution and a droplet was called to be positive for this HTO if it was predicted to belong to the component with the higher mean expression. Droplets positive for multiple HTOs were classified as doublets. Inter HTO doublets were found using the recoverDoublets function form the scDblFinder [45] package. Droplets negative for HTO library as well as doublets were filtered out. Droplets with < 500 UMIs, < 200 genes detected, and > 1% mitochondrial gene content were filtered out.

The decontaminated matrices were then imported into Seurat as individual objects, and SCTransform normalisation was performed for each library. We initially generated a preliminary low-resolution map of cell clusters for every sample, before running SoupX for ambient RNA removal [46]. PCA was performed on the corrected counts and a shared-nearest-neighbour graph built on these PCs (FindNeighbors) was clustered with the Louvain algorithm and a two-dimensional embedding was obtained with UMAP. The resulting cluster labels and UMAP coordinates, together with the raw UMI matrices, were then supplied to SoupX (SoupChannel, setClusters, setDR). For every sample, raw UMI counts together with the preliminary Seurat clusters and UMAP coordinates were loaded into SoupChannel. Background-corrected counts were generated with adjustCounts.

The ambient-RNA decontaminated matrices were then re-imported into Seurat as individual objects. Each dataset was normalised independently with SCTransform. Batch effects were regressed out with Harmony via IntegrateLayers. A shared nearest-neighbour graph was built on the Harmony space, and two-dimensional visualisation was obtained with RunUMAP. The resulting clusters and UMAP coordinates constituted the basis for subsequent analyses on the complete dataset. Cell types were annotated based on canonical gene markers [47; 48].

### 2.5. Single-nucleus RNA sequencing of human skeletal muscle

Muscle biopsies from the vastus lateralis of participants in the placebo group from a recently conducted clinical trial [49] were used for snRNA-seq analysis. In this trial, healthy adults (9 men and 6 women) were subjected to a protocol of eccentric contraction-induced muscle injury. Muscle biopsies were collected before, 2 hours after, and 2, 8, and 30 days after injury. Only muscle biopsies collected before the injury protocol were used. Nuclei isolation, snRNA-seq, data processing, clustering and cell type annotation were performed using the same methods used for mouse skeletal muscle tissue with adjustments specific to human tissue. RNA libraries were pseudoaligned to the GENCODE v44 reference transcriptome. Droplets with < 500 UMIs, < 200 genes, > 1% mitochondrial genes and < 15% counts from unspliced transcripts were filtered out.

### 2.6. AICAR and MK-8722 tolerance test

Mice were fasted for 3 hours (07:00–10:00 h) prior to the experiment. AICAR (250 mg/kg body weight) was injected intraperitoneally, and blood glucose levels were monitored for 120 min using the Contour XT glucometer (Hounisen, Denmark) and Contour Next glucose strips. MK-8722 tolerance was assessed by oral administration of either MK-8722 (10 mg/kg body weight) or vehicle (0.25% (w/v) methylcellulose, 5% (v/v) Polysorbate 80, and 0.02% (w/v) sodium lauryl sulfate in deionized water) [8; 30]. Blood glucose measurement was performed as described for the AICAR tolerance test.

### 2.7. Ex vivo skeletal muscle incubation and measurement of glucose uptake

Animals were anesthetized with Avertin [stock of 1 g/mL tribromoethanol (#T48402, Merck Millipore Sigma) in 2-methyl-2-butanol (#152463, Merck Millipore Sigma), diluted 1:20 in saline, and administered at 10 μL/g body weight via intraperitoneal injection. EDL muscles were rapidly dissected and mounted in oxygenated (95% O_2_ and 5% CO_2_), and warmed (30°C) Krebs-Ringer buffer (KRB) supplemented with 2 mM pyruvate in the presence of the indicated drug or vehicle for 50 min. The muscles were further incubated for 10 min in glucose uptake buffer (KRB buffer containing 1.5 μCi/mL [^3^H]-2-deoxy-D-glucose, 1 mM 2-deoxy-D-glucose, 0.45 μCi/mL [^14^C]-mannitol, 7 mM mannitol). At the end of the incubation period, muscles were frozen in liquid nitrogen and processed for glucose uptake assay [50] and immunoblot analysis.

### 2.5. Preparation of mouse tissue extracts for protein analysis

EDL muscles were homogenized in ice-cold lysis buffer (270 mM sucrose, 50 mM Tris-HCl (pH 7.5), 1 mM EDTA, 1 mM EGTA, 1% (v/v) Triton X-100, 20 mM glycerol-2-phosphate, 50 mM NaF, 5 mM Na_4_P2O_7_, 1 mM DTT, 0.1 mM PMSF, 1 mM benzamidine, 1 μg/mL microcystin-LR, 2 μg/mL leupeptin, and 2 μg/mL pepstatin A) using a TissueLyser II (Qiagen) at 30 Hz for 1 min. Gastrocnemius and quadriceps muscles were powdered and 10-15mg of the powdered tissue was used for lysis. Tissue lysates were centrifuged at 6,000 × g for 10 min at 4°C, and total protein concentration in the clarified lysate was determined by a microplate reader (Hidex Sense) using Bradford reagent (#23200, ThermoFisher) and BSA as standard.

### 2.8. Immunoblotting

Protein extracts were denatured in Laemmli buffer at 95°C for 5 min. Proteins (12-15 µg) were separated by SDS-PAGE on 4–20% gradient gels (Criterion™ TGX Stain-Free™, # 5678095, BioRad) and transferred onto nitrocellulose membranes (#926-31090, LI-COR). Membranes were blocked in 5% skimmed milk for 1 h at room temperature. The membranes were subsequently incubated in TBST (10 mM Tris-HCl (pH 7.6), 137 mM NaCl, and 0.1% (v/v) Tween-20) containing 5% (w/v) BSA and primary antibody overnight at 4°C. The membranes were extensively washed before incubation with horseradish peroxidase (HRP)-conjugated secondary antibodies diluted 1:10,000 for 45 min. Signal detection of the blots was performed using enhanced chemiluminescence substrate (Millipore, WBKLS0500) and protein bands were visualized by Odyssey® XF imaging system. The volume densities of the protein bands were quantified with the ImageStudio Lite Ver 5.2 software. Specific blot/band signals in each lane alongside housekeeping protein (vinculin) were obtained, and the background signal was subtracted.

### 2.9. Immunoprecipitation of AMPK and in vitro activity assay

Lysates of gastrocnemius muscle (100 μg) were incubated with a mix of 5-7.5 μL packed Protein G Sepharose beads and either polyclonal rabbit α1- or α2-AMPK antibody or IgG (as control) on an orbital shaker at 4°C for 2 hours (**Supplementary Table 1**). The beads were pelleted at 500 x g for 1 min and washed twice with 0.5 mL lysis buffer containing 500 mM NaCl and then washed twice with buffer A [50 mM HEPES (pH 7.4), 150 mM NaCl, 1 mM EGTA and 1 mM DTT]. Activity of the AMPK complexes were determined by phosphorylation of the synthetic peptide substrate AMARA (AMARAASAAALARRR) [51].

### 2.10. Statistical analysis

Data are expressed as mean ± SEM unless otherwise indicated. Statistical analyses were conducted using GraphPad Prism software version 10.5.0 (774) (La Jolla, CA, USA). Comparison between multiple groups was performed using two-way analysis of variance (ANOVA) followed by either Tukey’s post hoc test or Sidak’s multiple comparison tests, as indicated in the respective figure legends. For the drug tolerance tests, unpaired two-tailed Welsch’s t-test was used. p values of <0.05 were considered statistically significant.

## 3. RESULTS

### 3.1. Single-nucleus RNA-sequencing (snRNA-seq) reveals distinct expression patterns of AMPKγ isoforms in different cell types and myofiber-types within skeletal muscle tissue

AMPKγ1 is ubiquitously expressed in tissues, and γ1-containing AMPK complexes have been reported to predominate in both human and mouse skeletal muscle. However, these estimates are derived from immunoprecipitation and immunoblotting analyses of whole skeletal muscle tissue lysates, which do not provide cell-type-specific resolution. To address this limitation, we performed single-nucleus RNA sequencing (snRNA-seq) to map the expression patterns of AMPK isoforms across distinct cell populations within skeletal muscle tissues from both mice and humans. Recent advances in single-cell RNA sequencing (scRNA-seq), snRNA-seq, and single-cell mass cytometry, have substantially improved the characterization of cellular heterogeneity in skeletal muscle. These methods have enabled the identification of diverse myogenic populations, such as muscle stem cells, progenitor cells (myoblasts), and various mature fiber types (type I, IIa, IIx, and IIb), as well as non-myogenic cell types, including fibro/adipogenic progenitors (FAPs), immune cells, adipocytes, endothelial cells, tenocytes, and neural cells [47; 52; 53]. In this study, we applied snRNA-seq to both mouse (n = 6) and human (n = 6) skeletal muscle and generated over 14,480 and 11,530 single-nucleus transcriptomes, respectively (**Figure 1A**, **Figure 2A, Supplementary Table 3, Supplementary Table 4**). Unsupervised shared nearest neighbour (SNN) clustering partitioned mouse nuclei into 15 distinct cell types (clusters), including myogenic and non-myogenic cell types, based on their transcriptomic profiles (**Figure 1B**). To identify these clusters, we analyzed the normalized expression levels and frequency of canonical cell-type genes, naming them according to their distinct expression patterns (**Figure 1C**). A subclustering analysis restricted to mature myofibers, defined by myosin heavy chain expression, was conducted to quantify AMPK isoform expression in both mice (**Figure 1K**) and humans (**Figure 2K**). In mouse skeletal muscle, we observed that *Prkaa1* was ubiquitously expressed across diverse skeletal muscle cell populations, whereas *Prkaa2* shows abundant expression in mature myofibers (**Figure 1D, E and L**). *Prkab1* showed low expression, detected in only a few nuclei across clusters (**Figure 1F**), and notably absent in myofiber nuclei (**Figure 1L**). In contrast, *Prkab2* showed higher expression across multiple clusters (Figure 1G) including myofibers (**Figure 1 L**). *Prkag1* was expressed at relatively similar levels across the majority of cell types (**Figure 1H and L**). *Prkag2* exhibited moderate to high expression in immune cells, FAPs, NMJ cells, and muscle stem cells, while its expression was minimal in myofibers (**Figure 1I and L**). Consistent with previous findings [7; 8], *Prkag3* was almost exclusively expressed in glycolytic type IIb myofibers (**Figure 1J and L**).

**Figure 1.**
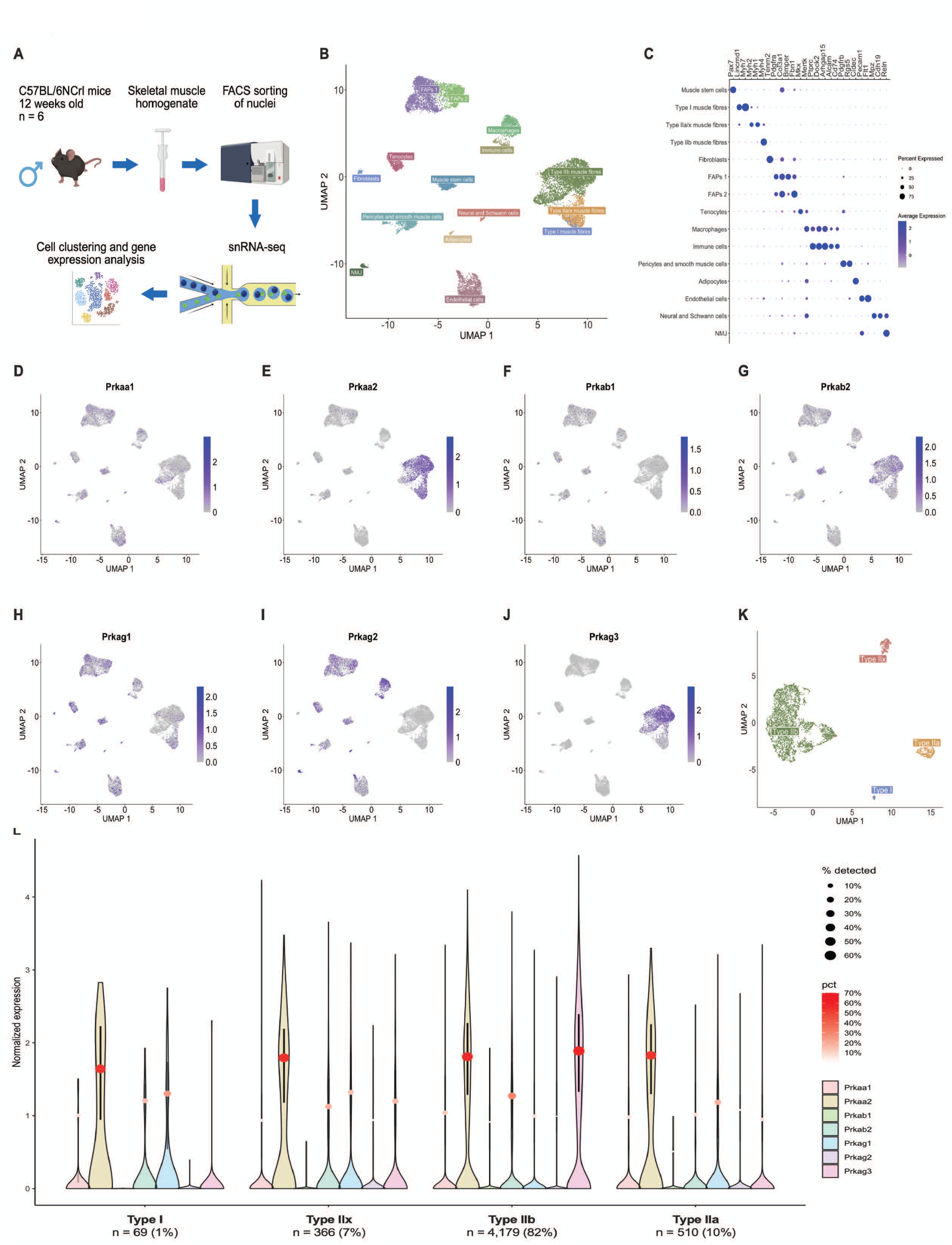
Gene expression patterns of AMPK subunit genes in mouse skeletal muscle. (A) Experimental design overview. The medial part of the mouse gastrocnemius muscle was used to isolate nuclei for single-nucleus RNA sequencing (snRNA-seq) and subsequent analyses of cell clustering and gene expression. (B) Complete snRNA-seq atlas. Over 14480 single-nucleus transcriptomes were generated and clustered into 15 distinct cell types, including myogenic and non-myogenic cells. Data are presented as a Uniform Manifold Approximation and Projection (UMAP). (C) Clusters were named based on the expression levels and frequency of canonical genes. Dot size indicates the percentage of nuclei with a non-zero expression level, while the colour scale reflects the average expression level across all nuclei in the cluster. (D-J) UMAPs illustrating the expression patterns of AMPK subunit genes, including *Prkaa1*, *Prkaa2*, *Prkab1*, *Prkab2*, *Prkag1*, *Prkag2*, and *Prkag3*. (K) UMAP visualization of myofibers (Type I, IIa, IIb, and IIx). (L) Violin plots showing normalized expression of AMPK subunit/isoform genes in myofibers. Dot and whisker represent median and interquartile range among expressing nuclei (expression > 0). Dot size and color indicate the percentage of nuclei with detectable expression.

**Figure 2.**
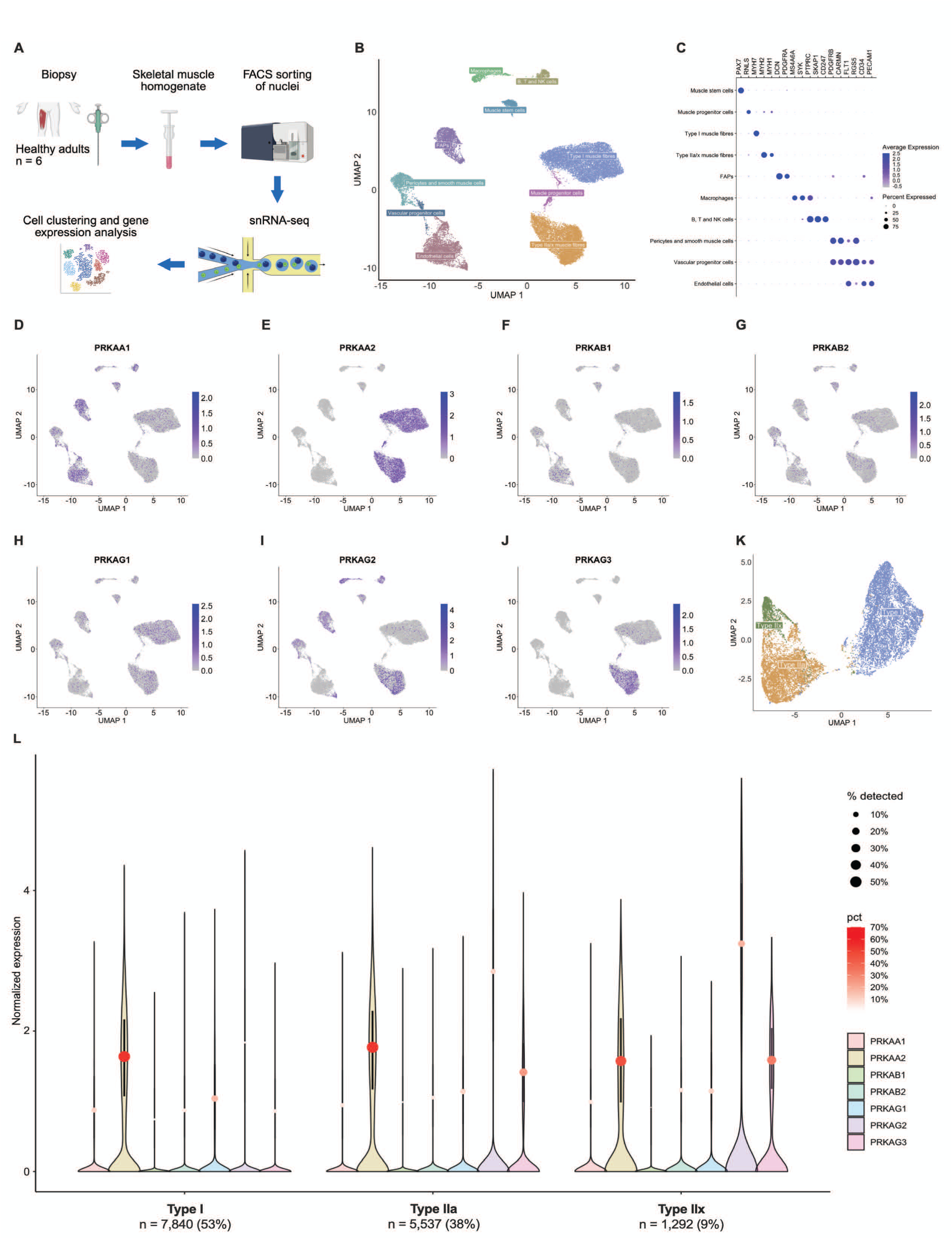
Gene expression patterns of AMPK subunit genes in human skeletal muscle. (A) Experimental design overview. Vastus lateralis muscle biopsies were used to isolate nuclei for snRNA-seq and subsequent analyses of cell clustering and gene expression. (B) Complete snRNA-seq atlas. Over 27694 single-nucleus transcriptomes were generated and clustered into 10 distinct cell types, including myogenic and non-myogenic cells. Data are presented as a UMAP projection. (C) Clusters were named based on the expression levels and frequency of canonical genes. Dot size indicates the percentage of nuclei with a non-zero expression level, while the colour scale reflects the average expression level across all nuclei in the cluster. (D-J) UMAPs illustrating the expression patterns of AMPK subunit genes, including *PRKAA1*, *PRKAA2*, *PRKAB1*, *PRKAB2*, *PRKAG1*, *PRKAG2*, and *PRKAG3*. (K) UMAP visualization of myofibers (Type I, IIa, and IIx). (L) Violin plots showing normalized expression of AMPK subunit genes in myofibers. Dot and whisker represent median and interquartile range among expressing nuclei (expression > 0). Dot size and color indicate the percentage of nuclei with detectable expression.

AMPK subunit isoform expression patterns in human skeletal muscle were largely similar to those observed in mice. SNN clustering identified 9 distinct clusters, including both myogenic and non-myogenic cell types (**Figure 2B and C**). *PRKAA1* was broadly but weakly expressed across all clusters (**Figure 2D**), whereas *PRKAA2* was predominantly expressed in mature myofibers (**Figure 2E and L**). Both *PRKAB1* and *PRKAB2* were sparsely expressed, appearing in only a few nuclei across multiple cell types (**Figure 2F, G and L**). Similarly, *PRKAG1* exhibited limited expression across clusters (**Figure 2H, I and L**). *PRKAG2* showed low-moderate expression in several clusters including pericytes, FAPs, macrophages and immune cells (**Figure 2I**). In contrast to the minimal *Prkag2* expression observed in mouse myofibers, *PRKAG2* showed slightly higher expression in human type IIa/IIx fibers (**Figure 2I and L**). *PRKAG3* expression was largely restricted to type IIa/IIx fibers (**Figure 2J and L**).

### 3.2. Time course of skeletal muscle-specific deletion of AMPKγ1 and γ3

As significant expression of AMPKγ2 was not detected at either the mRNA level (within nuclei) or protein level in mouse skeletal muscle fibers or myocytes (**Figure 1N and O**, [11]), we hypothesized that simultaneous deletion of AMPKγ1 and γ3 in skeletal muscle would abolish AMPK activator-stimulated glucose uptake. To test this hypothesis, we generated inducible, skeletal muscle-specific double AMPKγ1/γ3 knockout mice (imγ1^-/-^/γ3^-/-^) by breeding AMPKγ1/γ3 double foxed mice (AMPKγ1^fl/fl^ [10]/AMPKγ3^fl/fl^ [8]) with mice expressing a tamoxifen-inducible Cre-recombinase driven by the human skeletal actin promoter (HSA-MCM) [38]. To determine the optimal time point for maximal and consistent deletion of the AMPKγ1 and γ3 proteins in skeletal muscle, AMPKγ1^fl/fl^/AMPKγ3^fl/fl^ x HSA-MCM (imγ1^-/-^/γ3^-/-^) and AMPKγ1^fl/fl^/AMPKγ3^fl/fl^ (control) mice were treated with tamoxifen for three consecutive days, and hindlimb skeletal muscles were harvested either 3 or 6 weeks post-treatment (**Figure 3A**). Immunoblot analysis revealed a substantial deletion of AMPKγ1 (∼80%) and γ3 (∼95%) protein levels in gastrocnemius muscle from imγ1^-/-^/γ3^-/-^ compared to control mice at 3 weeks, with AMPKγ1 levels further reduced to >95% by 6 weeks (**Figure 3B**). The residual AMPKγ1 signal likely reflects expression in non-myogenic cell populations residing within the skeletal muscle tissue (**Figure 1**). Given the structural role of the γ subunit in stabilizing the AMPK heterotrimeric complex, deletion of γ1 and γ3 resulted in a marked reduction (∼50-80%) in the α- and β-subunits, with the most pronounced decreases observed for the α2 and β2 isoforms (**Figure 3B**). Similar results were observed in quadriceps muscle (**Supplementary Figure 1A and B**). Based on these findings, all subsequent biochemical and physiological experiments in control and imγ1^-/-^/γ3^-/-^ mice were conducted 6 weeks after tamoxifen administration.

**Figure 3.**
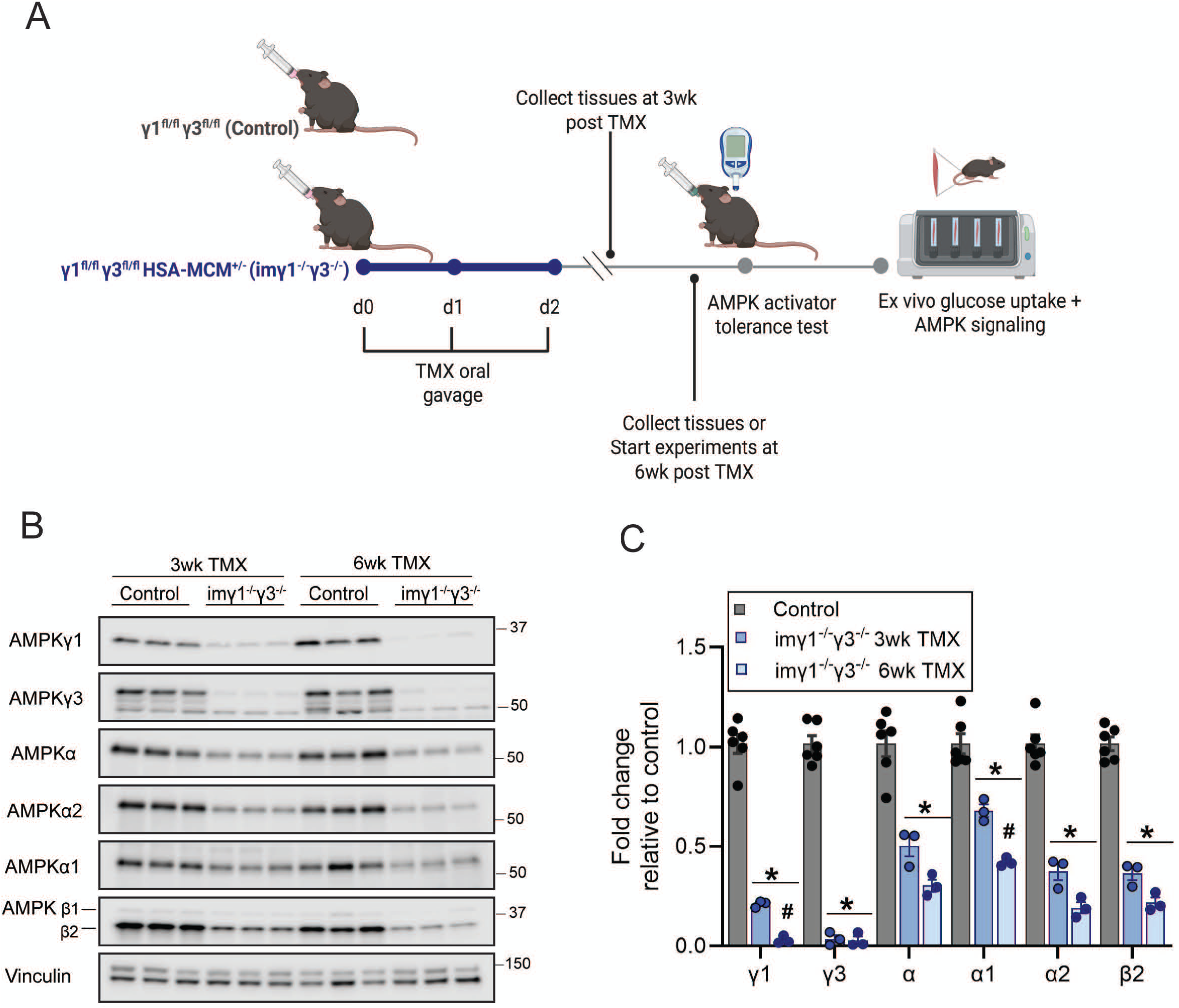
Tamoxifen-induced dual deletion of AMPKγ1 and γ3 in mouse skeletal muscle. (A) Schematic of the tamoxifen (TMX) regimen and experimental timeline. Skeletal muscle-specific deletion of AMPKγ1 and γ3 isoforms was achieved by expressing a TMX-inducible Cre-recombinase under the human α-skeletal actin promoter (HSA-MCM) (imγ1⁻^/^⁻/γ3⁻^/^⁻). Male mice (8-10 weeks old) from the indicated genotypes received an oral gavage of TMX (40 mg/kg body weight) for 3 consecutive days. The indicated experimental procedures were performed at either 3 weeks or 6 weeks after the final TMX dose. (B-C) Representative immunoblots and quantification of AMPK isoform expression in gastrocnemius muscle. Signals were normalized to vinculin (loading control) and are shown as fold change relative to control. Data are mean ± SEM; n = 6 per genotype. Statistical significance was assessed by two-way ANOVA with Sidak’s multiple comparisons test; \**P* < 0.05 (control γ1^fl/fl^/γ3^fl/fl^ *vs*. imγ1⁻^/^⁻/γ3⁻^/^⁻).

### 3.3. Genetic deletion of AMPKγ1/γ3 abolishes activator-induced but not insulin-stimulated glucose uptake in skeletal muscle *ex vivo*

To determine the roles of AMPKγ1 and γ3 in activator-mediated signaling and glucose uptake, we examined EDL muscles isolated from control and inducible imγ1⁻^/^⁻/γ3⁻^/^⁻ mice following *ex vivo* treatment with either vehicle or the AMP-mimetic AMPK activator, AICAR. In control muscles, AICAR significantly increased phosphorylation of AMPKα and its canonical substrate acetyl-CoA carboxylase 2 (ACC2) (**Figure 4A-D**). Given that ACC2 phosphorylation reaches saturation rapidly upon AMPK activation, we also assessed phosphorylation of starch binding domain-containing protein 1 (STBD1), a substrate with greater dynamic range [54]. In control muscles, AICAR treatment elicited a pronounced increase in STBD1 phosphorylation (**Figure 4A and E**). In contrast, AICAR failed to activate AMPK in imγ1⁻^/^⁻/γ3⁻^/^⁻ muscles, as evidenced by unchanged p-AMPKα/total AMPKα ratios. Although slight increases in ACC2 and STBD1 phosphorylation were detected in imγ1⁻^/^⁻/γ3⁻^/^⁻ muscles, these responses were markedly attenuated relative to controls (**Figure 4A, D, and E**). Consistent with these findings, AICAR-stimulated glucose uptake was robustly increased (∼2.5-fold increase) in control muscles but was abolished in imγ1⁻^/^⁻/γ3⁻^/^⁻ muscles (**Figure 4F**). In contrast, proximal insulin signaling and insulin-stimulated glucose uptake remained intact in imγ1⁻^/^⁻/γ3⁻^/^⁻ muscles (**Figure 4G, Supplementary Figure 1C**). These results demonstrate that the deletion of AMPKγ1/γ3 selectively impairs AICAR-induced but not insulin-mediated (*i.e.*, AMPK independent) glucose uptake.

**Figure 4.**
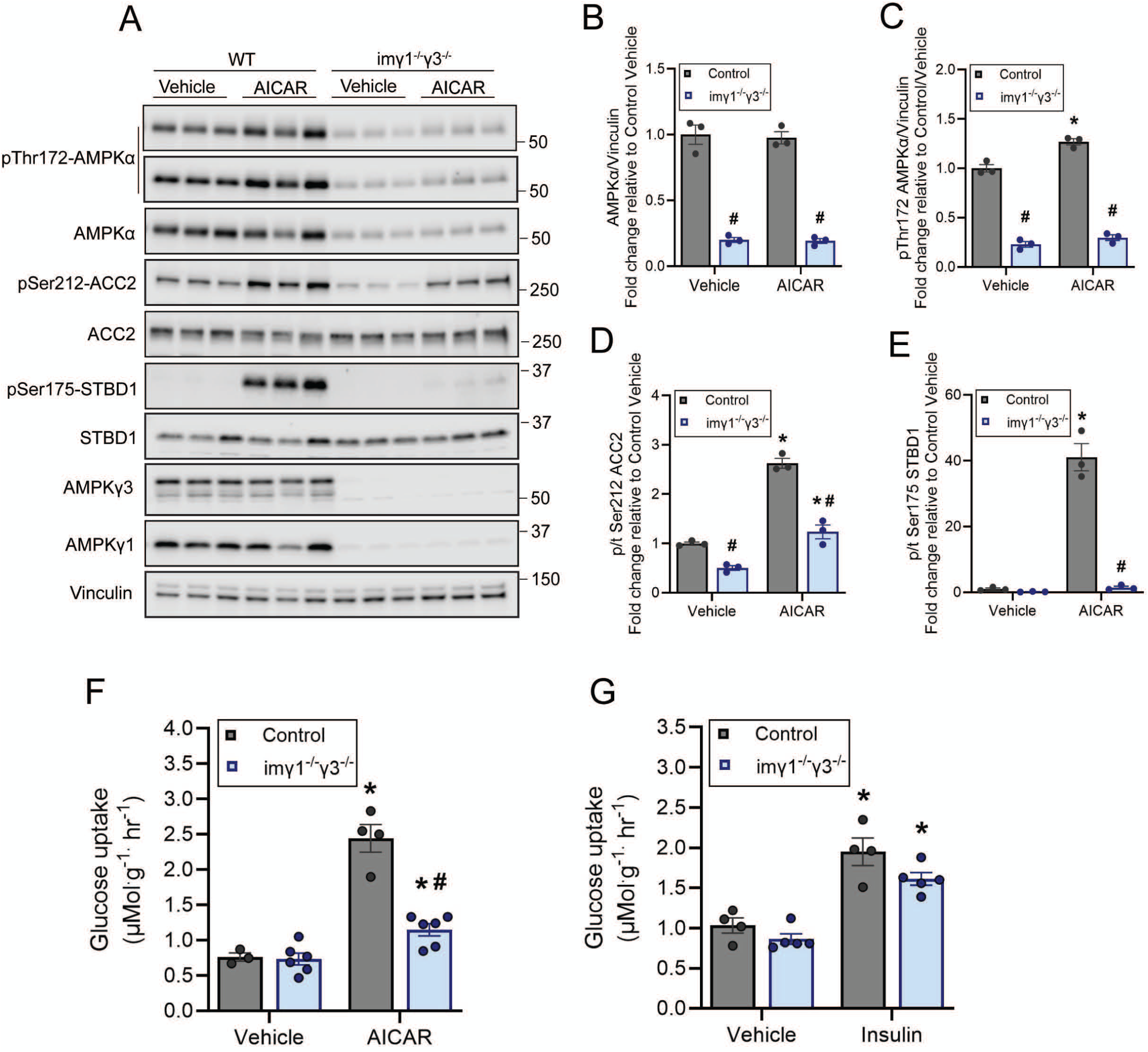
Skeletal muscle-specific deletion of AMPKγ1 and γ3 abolishes AICAR-induced, but not insulin-stimulated, glucose uptake. (A–F) Six weeks after tamoxifen (TMX) treatment, extensor digitorum longus (EDL) muscles from the indicated genotypes were incubated with vehicle (0.1% DMSO) or AICAR (2 mM) for 50 min, followed by a 10-min incubation with radioactive 2-deoxyglucose. (A–E) Portions of the muscle extracts were analyzed by immunoblotting with the indicated antibodies; (F) the remainder was used for glucose-uptake measurements (n = 4-6 per treatment/genotype). Data are representative of two independent experiments. (G) EDL muscles from control γ1^fl/fl^/γ3^fl/fl^ and imγ1⁻^/^⁻/γ3⁻^/^⁻ mice, after 6 weeks after TMX treatment, were incubated with or without insulin (100 nM) for 50 min followed by a 10-min incubation with radioactive 2-deoxyglucose followed by glucose uptake measurement (n= 4-5 per treatment/genotype). Results are representative of two independent experiments. Data are shown as mean ± SEM. Statistical significance was determined by two-way ANOVA with Tukey’s post hoc correction. \**P* < 0.05 (treatment effect within the same genotype), ^#^*P* < 0.05 (genotype effect within the same treatment).

We next assessed the effect of MK-8722, an ADaM site activator [30], on glucose uptake in control and imγ1⁻^/^⁻/γ3⁻^/^⁻ EDL muscles. Consistent with a previous study and similar to the effect of AICAR (**Figure 4A-E**), MK-8722 significantly increased phosphorylation of AMPKα, ACC2 and STBD1 in control [8]. In contrast, these phosphorylation responses were markedly attenuated in imγ1⁻^/^⁻/γ3⁻^/^⁻ muscles (**Figure 5A-E**). Likewise, MK-8722-stimulated glucose uptake *ex vivo* was fully abolished in the imγ1⁻^/^⁻/γ3⁻^/^⁻ muscles (**Figure 5F**). Supporting these *ex vivo* findings, blood glucose lowering in response to MK-8722 administration *in vivo* was significantly blunted in imγ1⁻^/^⁻/γ3⁻^/^⁻ mice compared to controls (**Figure 5G**). Collectively, these results demonstrate that γ1 and γ3, but not γ2, are essential for AMPK activator-stimulated glucose uptake in mouse skeletal muscle.

**Figure 5.**
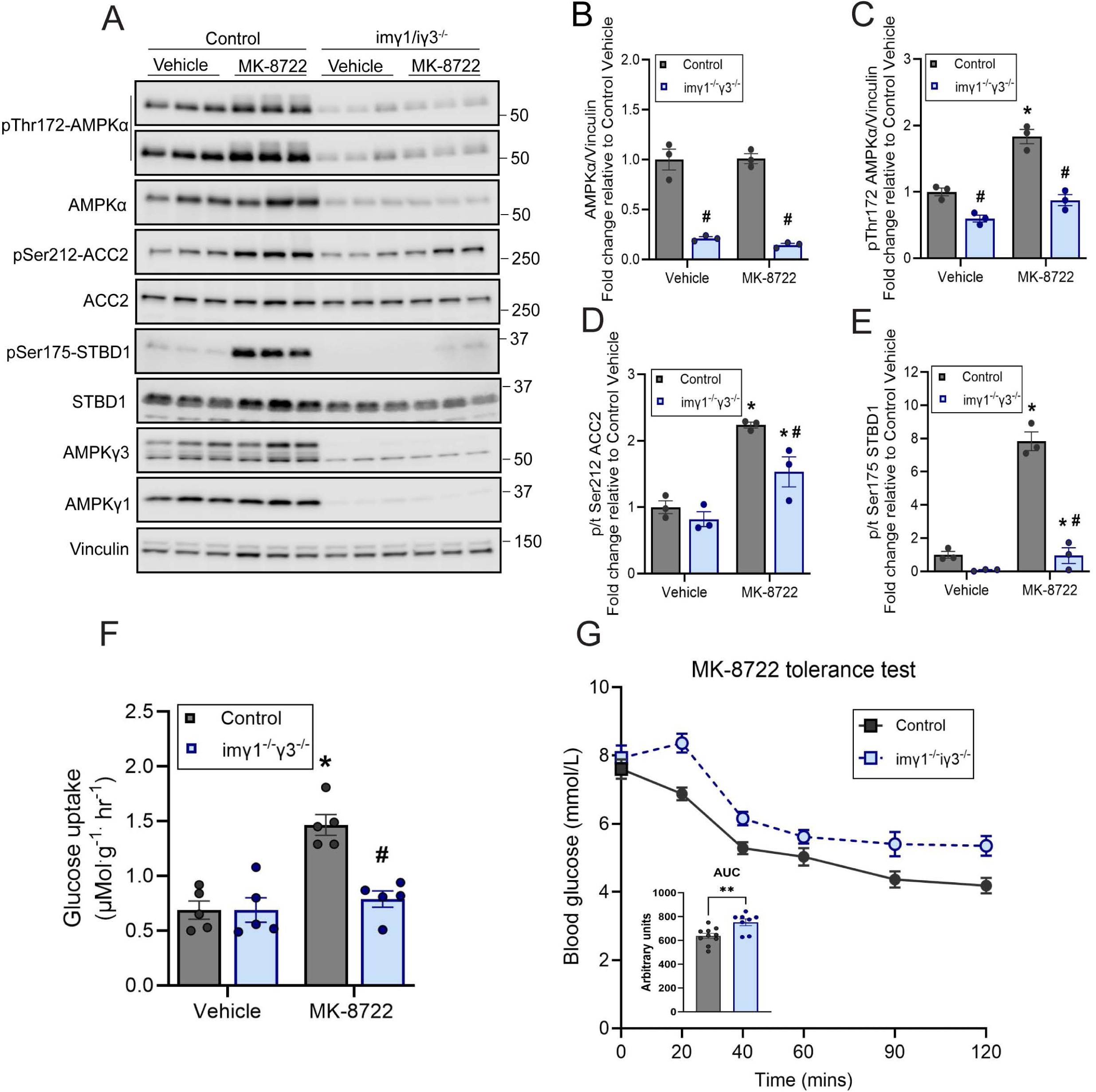
Skeletal muscle-specific deletion of AMPKγ1 and γ3 impairs MK-8722-induced muscle glucose uptake *ex vivo* and blood glucose lowering *in vivo*. (A–F) Six weeks after tamoxifen (TMX) treatment, extensor digitorum longus (EDL) muscles from the indicated genotypes were incubated with vehicle (0.1% DMSO) or MK-8722 (10 µM) for 50 min, followed by a 10-min incubation with radioactive 2-deoxyglucose. (A–E) Portions of the muscle extracts were analyzed by immunoblotting with the indicated antibodies; (F) the remainder was used for glucose-uptake measurements (n = 5 per treatment/genotype). Data are representative of two independent experiments. (G) MK-8722 tolerance test: imγ1⁻^/^⁻/γ3⁻^/^⁻ mice were fasted for 3 h, orally gavaged with MK-8722 (10 mg/kg), and blood glucose was monitored for the indicated time. Area under the curve (AUC) was calculated for each group (n = 8 per genotype). Data are mean ± SEM. \**P* < 0.05 (treatment effect within the same genotype), ^#^*P* < 0.05 (genotype effect within the same treatment). Statistical significance was determined by two-way ANOVA with Tukey’s post-hoc correction (A–F) or unpaired two-tailed Welch’s t-test (G).

### 3.4. Skeletal muscle-specific deletion of γ1 does not impair AMPK activator-induced muscle glucose uptake *ex vivo* or blood glucose lowering *in vivo*

We and others previously demonstrated that loss of AMPKγ3 selectively impairs AICAR-stimulated, but not ADaM site activator-stimulated, glucose uptake in skeletal muscle [8; 36; 37]. Given that the γ1-containing AMPK complex (α2β2γ1) is the predominant isoform in skeletal muscle [5; 6], we sought to determine if γ1 is required for ADaM site activator-stimulated glucose uptake. To this end, inducible, skeletal muscle-specific AMPKγ1⁻^/^⁻ mice (imγ1^-/-^) were generated by crossing AMPKγ1^fl/fl^ mice with mice expressing a tamoxifen-inducible Cre-recombinase under control of the HSA-MCM promoter. iImγ1^-/-^ and control (AMPKγ1^fl/fl^) mice received tamoxifen treatment for three consecutive days, and hindlimb skeletal muscles were harvested six weeks post-treatment (**Figure 6A**) for analyses of AMPK subunit/isoform expression, *ex vivo* glucose uptake, and AMPK signaling responses to pharmacological activators. As expected, based on the findings in imγ1⁻^/^⁻/γ3⁻^/^⁻ mice, deletion of γ1 alone resulted in near-complete ablation of γ1 expression and activity without altering γ3 levels or activity in gastrocnemius (**Figure 6B-D**) and quadriceps muscles (**Supplementary Figure 1D and E**). Loss of γ1 also led to an approximately 50% reduction in α1/α2 and β2 expression (**Figure 6C and D, Supplementary Figure 1D and E**). AICAR-stimulated AMPK activity, as assessed via p-AMPKα (normalized to vinculin, as total AMPKα was profoundly reduced) and phosphorylation of ACC2 and STBD1, was comparable between imγ1^-/-^ and control EDL muscles (**Figure 7A-E**). Correspondingly, AICAR-stimulated glucose uptake in EDL muscle *ex vivo* (**Figure 7F**) and AICAR-induced blood glucose lowering and recovery kinetics *in vivo* were similar between groups (**Figure 7G**). As a control, skeletal muscle-specific inducible AMPKγ3⁻^/^⁻ mice (imγ3^-/-^) were generated and analyzed (**Supplementary Figure 2A**). Consistent with our previous findings using constitutive AMPKγ3⁻^/^⁻ mice [8], inducible γ3 deletion led to a 20-30% reduction in AMPKα expression (**Supplementary Figure 2B, C, F and G**) and a marked loss of AICAR-, but not MK-8722-stimulated glucose uptake and corresponding AMPK phosphorylation in EDL muscle *ex vivo* (**Supplementary Figure 2B-I**).

**Figure 6.**
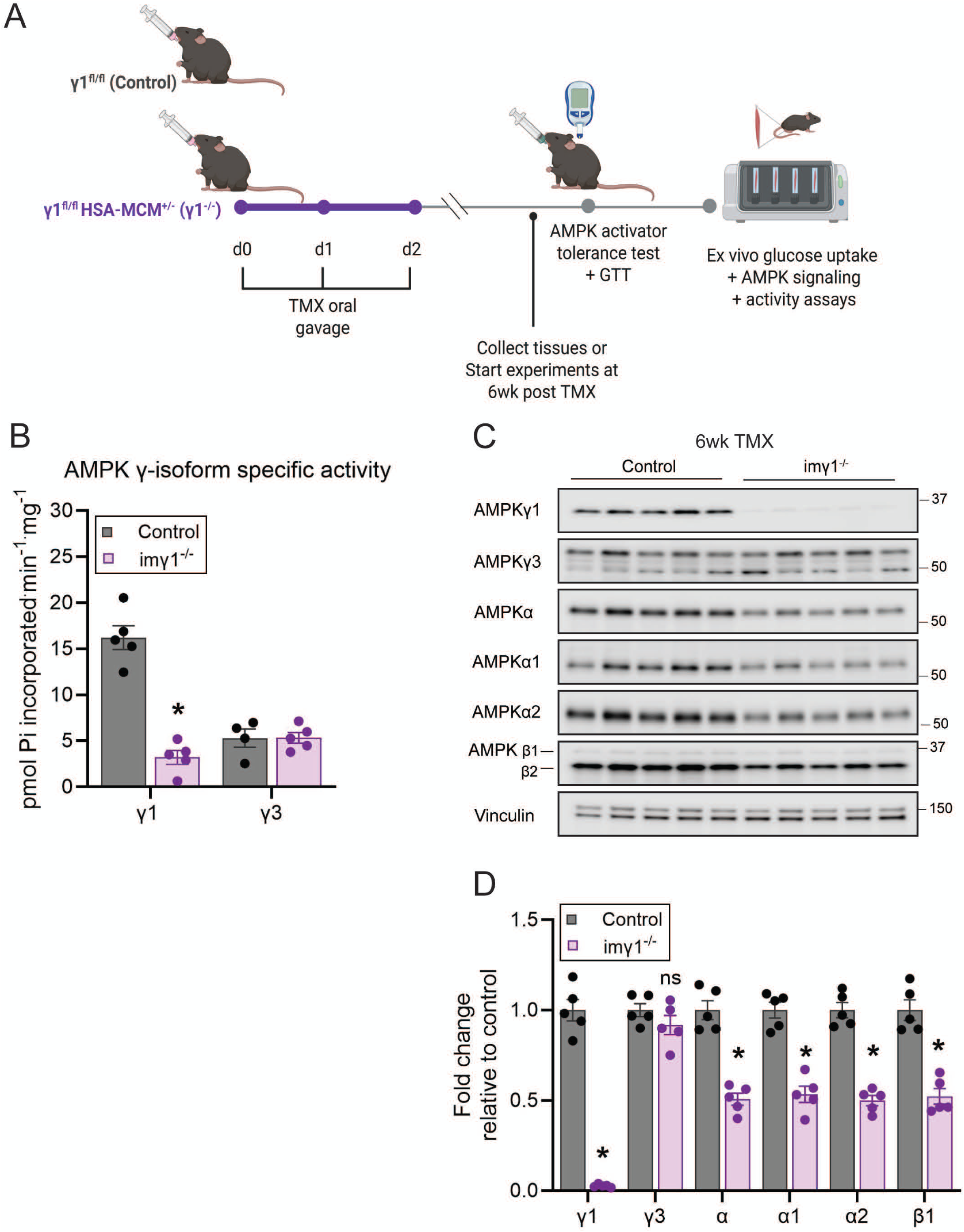
Tamoxifen-induced skeletal muscle deletion of AMPKγ1. (A) Schematic of the tamoxifen (TMX) regimen and experimental timeline. Skeletal muscle-specific deletion of AMPKγ1 was achieved by expressing a TMX-inducible Cre-recombinase under the human α-skeletal actin promoter (HSA-MCM) (imγ1⁻^/^⁻). Male mice (8-10 weeks old) from the indicated genotypes received an oral gavage of TMX (40 mg/kg body weight) for 3 consecutive days. Experiments were performed 6 weeks post TMX treatment. (B) The γ1- and γ3-containing AMPK complexes were immunoprecipitated from gastrocnemius muscles harvested from the indicated genotypes and an *in vitro* AMPK activity assay was performed in duplicate (n = 5 per genotype). (C-D) Representative immunoblots and quantification of AMPK isoform expression in gastrocnemius muscle. AMPK isoform expressions were normalized to vinculin expression (loading control) and are shown as fold-change relative to control (n = 8 per treatment/genotype). Data are shown as mean ± SEM. Statistical significance was determined using two-way ANOVA with Sidak’s multiple comparisons test and are shown as \**P* < 0.05 (WT *vs*. iγ1^-/-^) (D). Statistical significance was determined using two-way ANOVA with Tukey’s post hoc correction (B) and are shown as \**P* < 0.05 (control *vs*. imγ1^-/-^).

**Figure 7.**
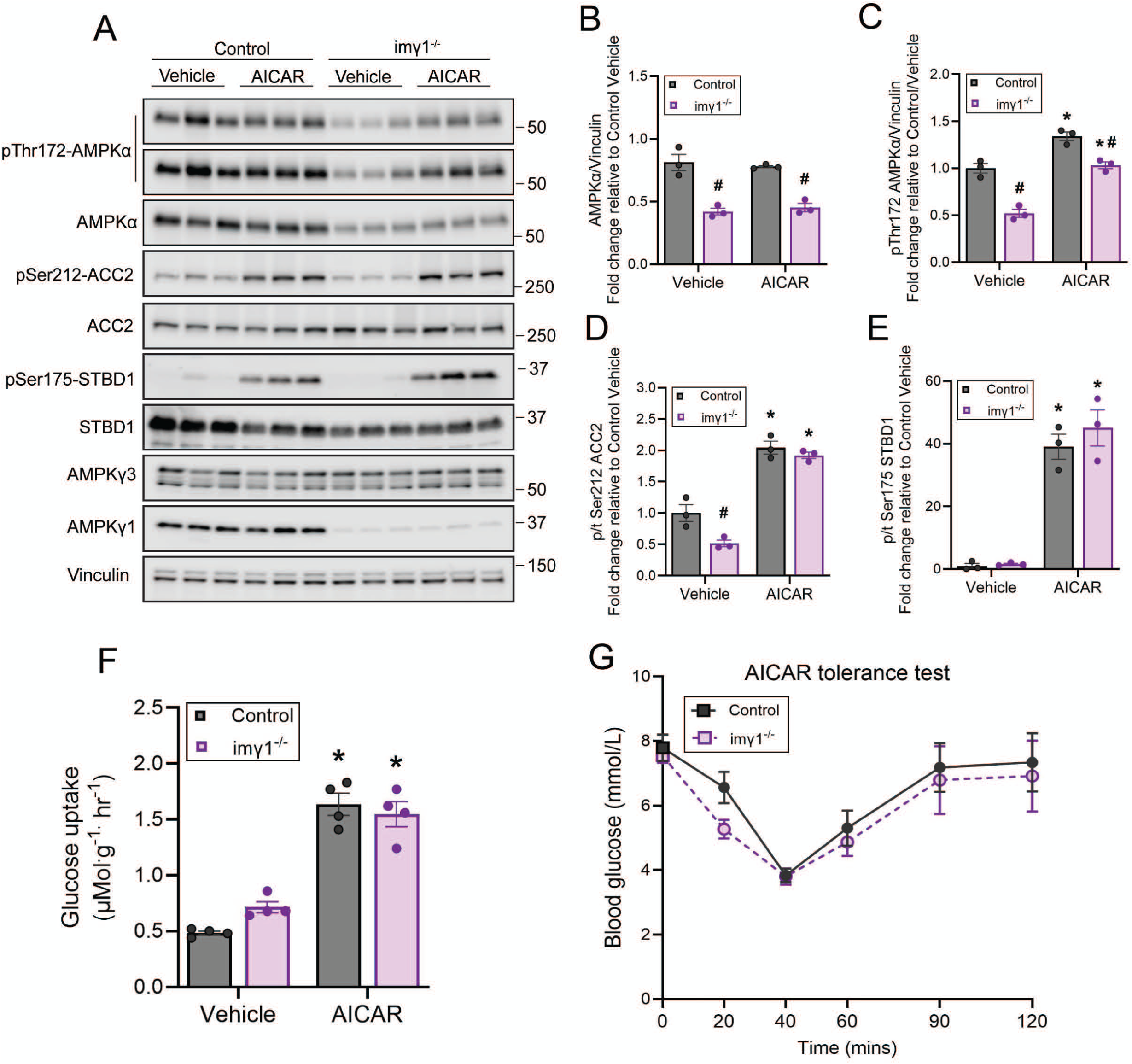
Skeletal muscle-specific deletion of AMPKγ1 does not alter AICAR-induced muscle glucose uptake *ex vivo* or blood glucose lowering *in vivo.* (A–F) Six weeks after tamoxifen (TMX) treatment, extensor digitorum longus (EDL) muscles from the indicated genotypes were incubated with vehicle (0.1% DMSO) or AICAR (2 mM) for 50 min, followed by a 10-min incubation with radioactive 2-deoxyglucose. (A–E) Portions of the muscle extracts were analyzed by immunoblotting with the indicated antibodies; (F) the remainder was used for glucose-uptake measurements (n = 4 per treatment/genotype). Data are representative of two independent experiments. (G) AICAR tolerance test: six weeks after TMX treatment, male imγ1⁻/⁻ mice were fasted for 3 h and intraperitoneally injected with AICAR (250 mg/kg body weight), followed by blood glucose monitoring over the indicated duration (n = 8 per genotype). Data are shown as mean ± SEM. Statistical significance was determined by two-way ANOVA with Tukey’s post hoc correction (A-F) or unpaired, two-tailed Welsch’s t-test for AUC (data not shown) using data in (G). \**P* < 0.05 (treatment effect within the same genotype), ^#^*P* < 0.05 (genotype effect within the same treatment).

In imγ1⁻^/^⁻ EDL muscle, while effects of MK-8722 on net p-AMPK and STBD1 phosphorylation were blunted (**Figure 8A-C and E**), MK-8722-induced ACC2 phosphorylation (**Figure 8A and D**), glucose uptake (**Figure 8F**), and blood glucose-lowering kinetics (**Figure 8G**) remained comparable to those observed in control mice.

**Figure 8.**
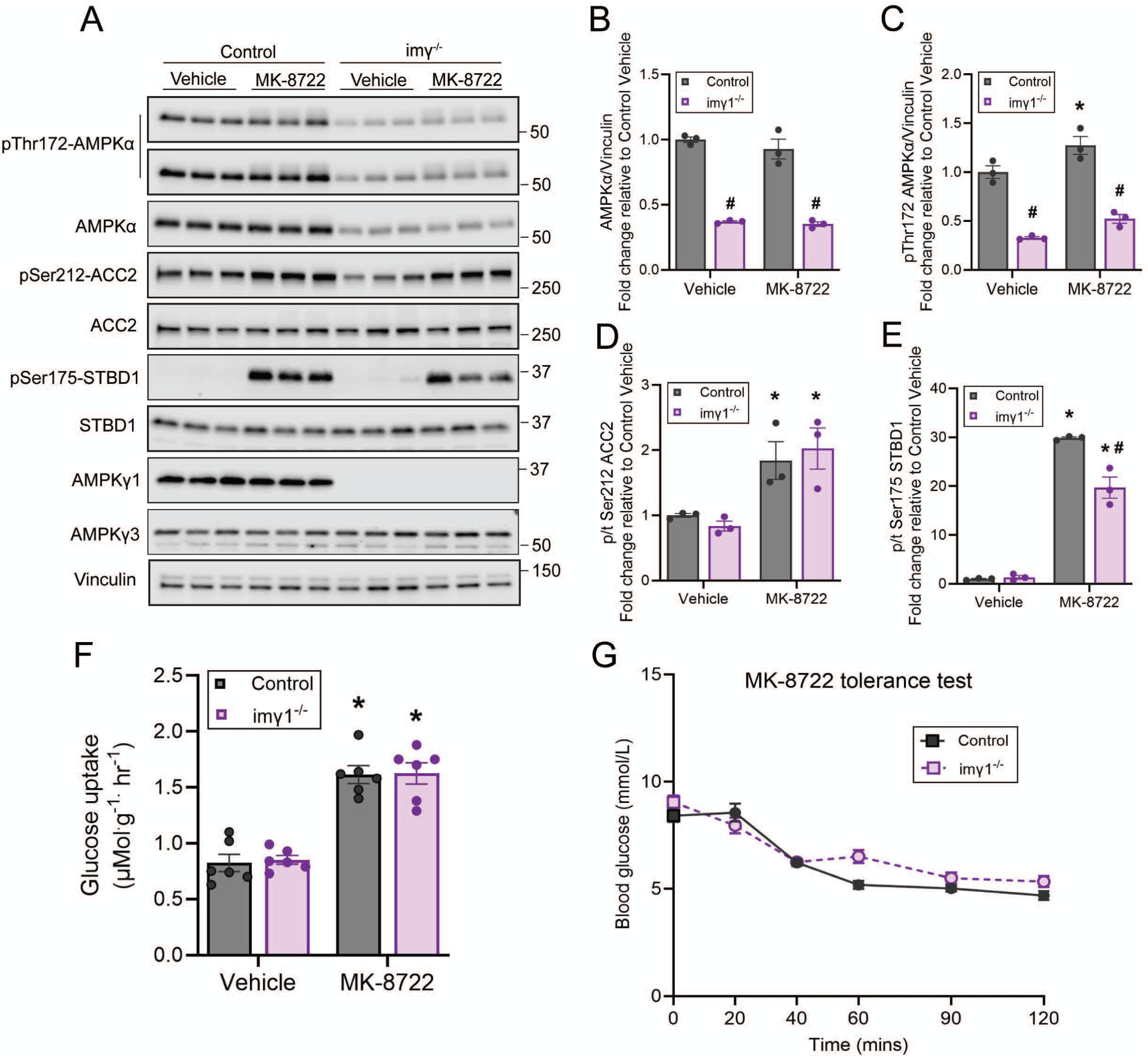
Skeletal muscle-specific deletion of AMPK γ1 does not alter MK-8722-induced muscle glucose uptake *ex vivo* or blood glucose lowering *in vivo.* (A–F) Six weeks after tamoxifen (TMX) treatment, extensor digitorum longus (EDL) muscles from the indicated genotypes were incubated with vehicle (0.1% DMSO) or MK-8722 (10 µM) for 50 min, followed by a 10-min incubation with radioactive 2-deoxyglucose. (A–E) Portions of the muscle extracts were analyzed by immunoblotting with the indicated antibodies; (F) the remainder was used for glucose-uptake measurements (n = 6 per treatment/genotype). Data are representative of two independent experiments. (G) MK-8722 tolerance test: imγ1⁻^/^⁻ mice were fasted for 3 h, orally gavaged with MK-8722 (10 mg/kg), and blood glucose was monitored for the indicated time (n = 8 per genotype). Data are mean ± SEM. Statistical significance was determined by two-way ANOVA with Tukey’s post hoc correction (A-F) or unpaired, two-tailed Welsch’s t-test for AUC (data not shown) using data in (G). \**P* < 0.05 (treatment effect within the same genotype), ^#^*P* < 0.05 (genotype effect within the same treatment).

Taken together, while γ1 is the predominant isoform stabilizing the AMPK trimers, it is dispensable for both AICAR-and MK-8722-induced glucose uptake in skeletal muscle.

## 4. DISCUSSION

The diverse biological functions of AMPK are thought to arise from the cell type- and tissue-specific expression, as well as subcellular localization, of its heterotrimeric complexes. Despite extensive investigation of AMPK signaling, the precise tissue-and subcellular-level concentrations of AMPK heterotrimers remain poorly defined. Among these, the AMPKα2β2γ1 complex has been reported to account for approximately 65-70% of the total AMPK trimers in human vastus lateralis and mouse EDL muscle [5; 6]. However, these estimates are based on immunoprecipitation and immunoblotting analyses of whole skeletal muscle lysates, which lack resolution at the cell-type level. Immunoblotting is inherently semi-quantitative, and accurate comparisons of AMPK subunit expression are constrained by variations in antibody specificity and sensitivity. In addition, detection of isoforms and splice variants may be incomplete, as single antibodies can fail to recognize epitopes that are absent, post-translationally modified, or conformationally masked, particularly in immunoprecipitation settings. Addressing this knowledge gap will require absolute quantification of AMPK subunits [55], ideally at single-myofiber resolution and across species, to more precisely define the molecular landscape of AMPK complexes in human skeletal muscle. Here, snRNA-seq analysis revealed distinct AMPK isoform expression patterns within skeletal muscle tissue in both humans and mice. *PRKAG1*, encoding AMPKγ1, was broadly expressed across multiple cell types, whereas *PRKAG3*, encoding AMPKγ3, was specifically enriched in fast twitch/glycolytic myofibers. Notably, *PRKAG2*, which encodes AMPKγ2, was sparsely expressed in mouse myogenic nuclei but was expressed in type IIa/IIx human myofibers. This pattern aligns with prior findings in primary human myotubes [7]) and suggests a potential species-specific difference in the expression of γ2 subunit. Interestingly, a recent phosphoproteomics study in human skeletal muscle identified a phosphorylation site on the γ2 subunit of AMPK (Ser196) that correlated with insulin-stimulated leg glucose uptake [56]. However, it remains unclear whether this γ2 regulation is intrinsic to myofibers or originates from non-muscle cells.

Consistent with our current snRNA-seq data and previous immunoblotting findings [11], which demonstrate that AMPKγ2 expression is minimal or undetectable in mouse skeletal muscle fibers, the simultaneous deletion of γ1 and γ3 led to a profound reduction in AMPK substrate phosphorylation and abrogation of glucose uptake in response to AMPK activator treatments *ex vivo*. This was accompanied by an ∼80% decrease in the expression of α and β subunits. As shown in the current and previous [57] studies given that subsets of total nuclei are reported to originate from non-myocyte populations, the residual expression of α and β subunits likely reflect AMPK trimer formation involving γ1 or γ2 isoforms in non-muscle cells. To date, there is no compelling evidence supporting the existence of stable αβ dimers in myofibers.

Notably, although deletion of γ1 led to an ∼50% reduction in α and β subunit expression without compensatory upregulation of γ3, both AICAR- and MK-8722–stimulated AMPK activation and glucose uptake remained fully preserved. These findings indicate that a reduced pool (∼50%) of γ3-containing AMPK trimers is sufficient to sustain full activation of AMPK and promote ADaM site activator-induced glucose uptake, while γ3 is specifically required for AICAR-stimulated glucose uptake in mouse skeletal muscle. However, the molecular mechanisms by which γ3-containing AMPK trimers selectively mediate AMP/ZMP-stimulated glucose uptake remain to be fully elucidated. All γ isoforms share a highly conserved C-terminal region containing the four cystathionine β-synthase domains responsible for nucleotide binding. Notably, γ2 and γ3 isoforms possess long N-terminal extensions absent in γ1. These extensions are intrinsically disordered and lack sequence conservation between isoforms, and their functional relevance remains unknown. To date, no high-resolution structures (crystal or cryo-EM) have been resolved for AMPK heterotrimers containing γ2 or γ3, leaving the structural contributions of these N-terminal regions unexplored. Future studies could benefit from the generation of γ3 knock-in mice lacking the N-terminal region to directly assess its physiological role. Additionally, spatial proteomics approaches in skeletal muscle would help delineate isoform-specific subcellular localization patterns of AMPKγ isoforms, offering further insight into their distinct regulatory functions.

## 5. CONCLUSIONS

While γ1 predominates in stabilizing the AMPKα2β2γ1 complex, it is dispensable for AMPK activator-stimulated glucose uptake in skeletal muscle, whether mediated via the nucleotide-binding or ADaM site.

## AUTHOR CONTRIBUTIONS

Conceptualization: K.S. Experimental design: D.B., E.E.G., C.J., J.B.J., J.F., J.G., J.T.T., M.A., K.S. Experimental execution: D.B., E.E.G., D.A., J.F., C.J., J.B.J., J.G., M.A. Data analysis: D.B., E.E.G., A.N.S., J.J., M.A. Data visualization and figure formatting: D.B., EEG, ANS, JJ. Resource/tool development: M.F. Supervision: JF, JJ, J.T.T., M.A., K.S. Writing-Original draft preparation: K.S. Writing-Reviewing and editing: All authors.

## GRANTS

The work is supported by the Novo Nordisk Foundation (NNF21OC0070257) to K.S. Novo Nordisk Foundation Center for Basic Metabolic Research is an independent Research Center based at the University of Copenhagen, Denmark, and partially funded by an unconditional donation from the Novo Nordisk Foundation (Grant number NNF23SA0084103).

## ACKNOWLEDGMENTS

We thank Karyn A. Esser (University of Florida) for her generous donation of the HSA-MCM mice. We acknowledge The Single-Cell Omics platform at the Novo Nordisk Foundation Center for Basic Metabolic Research (CBMR) for the technical expertise and support. We thank Kaltum Mohammed and Luna Krusholt for their technical support with genotyping. Schematic illustrations were created with BioRender.com.

## CONFLICT OF INTEREST

The authors declare that there are no competing interests associated with the manuscript.

## DATA AVAILABILITY

RNA-seq data pertaining to this work has been deposited in the Gene Expression Omnibus (GEO) repository under accession GSE306977. Reviewer access token (GEO): ulefcycodluhlcn

---

**Supplementary Figure 1.**
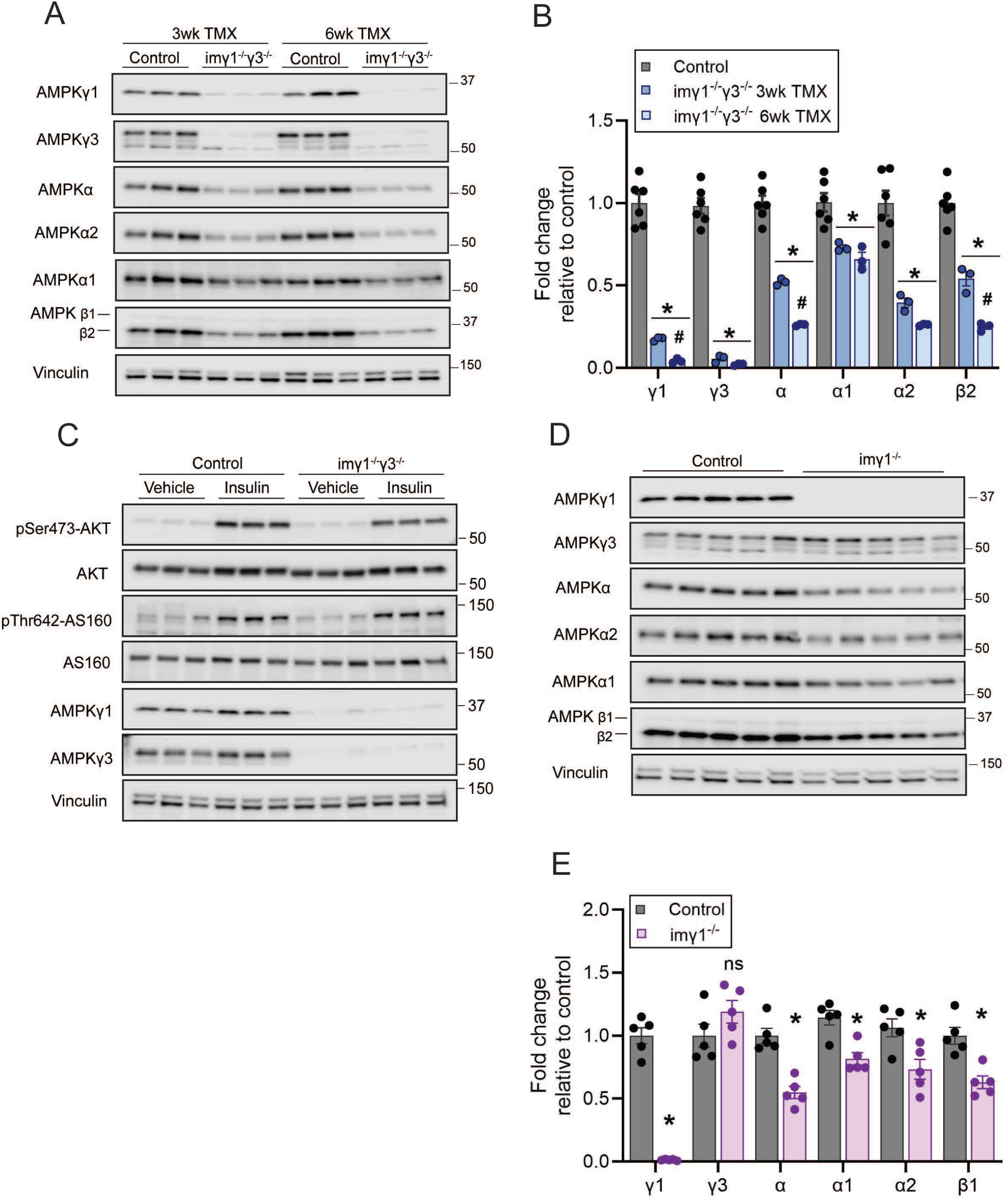
(A-B) Representative immunoblots and quantification of AMPK isoform expression in the quadriceps muscles from imγ1⁻^/^⁻/γ3⁻^/^⁻ mice. AMPK isoform expression was normalized to vinculin and is shown as fold change relative to control. Results are representative of two independent experiments performed with tissues from 8 mice (per genotype). Data are shown as mean ± SEM. Statistical significance was determined by two-way ANOVA with Sidak’s multiple comparisons test and are shown as \**P* < 0.05 (control *vs*. iγ1γ3-/-). n = 6 per group/genotype (C) EDL muscles from control γ1^fl/fl^/γ3^fl/fl^ and imγ1⁻^/^⁻/γ3⁻^/^⁻ mice, after 6 weeks after TMX treatment, were incubated with or without insulin (100 nM) for 50 min followed by a 10-min incubation with radioactive 2-deoxyglucose. Immunoblot analysis was performed using the indicated antibodies (n = 4-5 per treatment/genotype). (D-E) Representative immunoblots and quantification of AMPK isoform expression in quadriceps muscles from imγ1^-/-^ mice. AMPK isoform expression was normalized to vinculin and is shown as fold change relative to control. Results are representative of two independent experiments performed with tissues from 8 mice. Data are shown as mean ± SEM. Statistical significance was determined by two-way ANOVA with Sidak’s multiple comparisons test and are shown as \**P* < 0.05 (control *vs*. iγ1^-/-^).

**Supplementary Figure 2.**
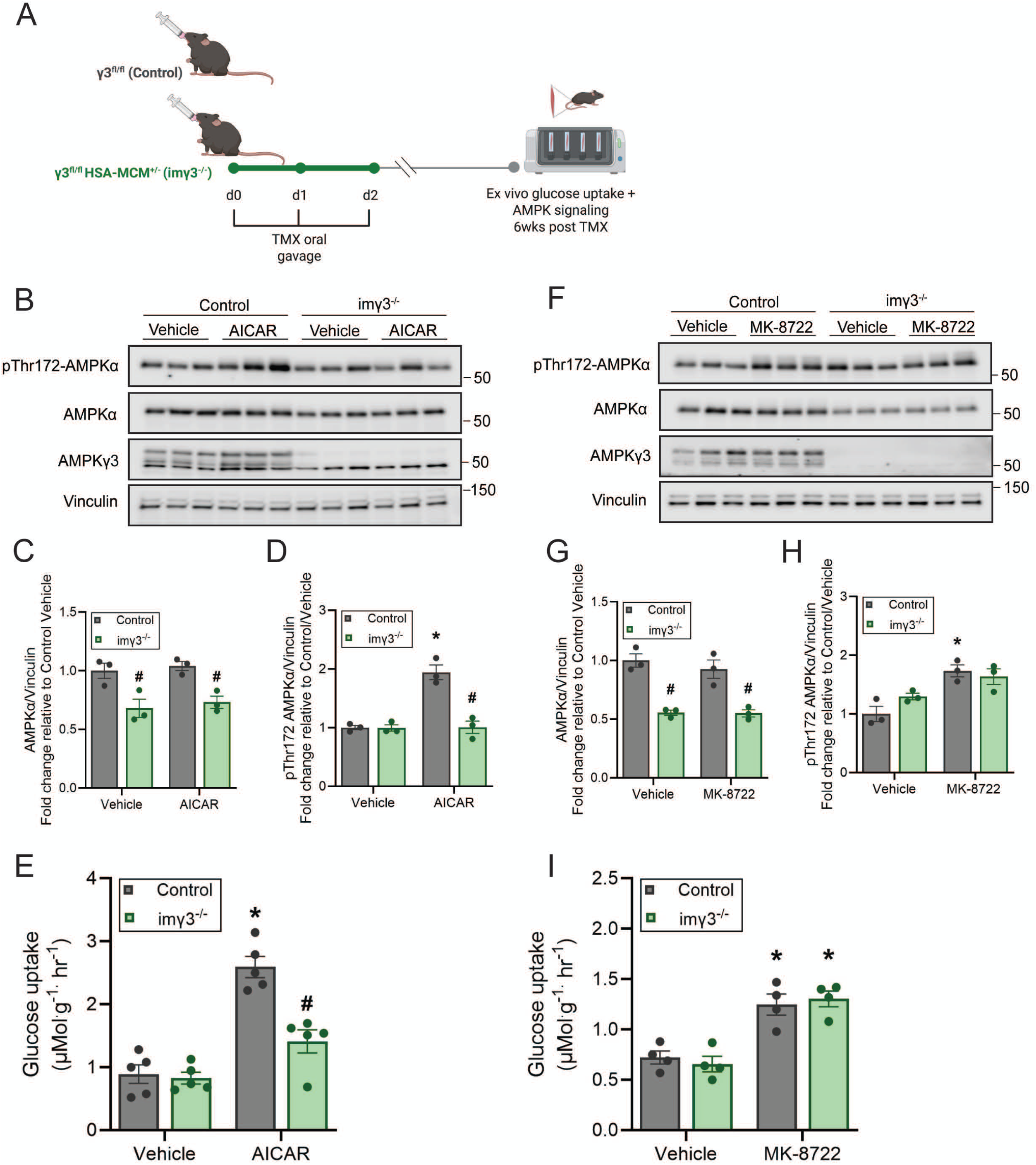
Skeletal muscle-specific TMX-inducible deletion of AMPK γ3 impairs AICAR-stimulated, but not MK-8722-stimulated glucose uptake. (A) Schematic of the tamoxifen (TMX) regimen and experimental timeline. Male mice (8-10 weeks old) from the indicated genotypes received an oral gavage of TMX (40 mg/kg body weight) for 3 consecutive days. Experiments were performed 6 weeks post TMX treatment. (B-I) Six weeks after tamoxifen (TMX) treatment, extensor digitorum longus (EDL) muscles from the indicated genotypes were incubated with vehicle (0.1% DMSO), AICAR (2 mM) or MK-8722 (10 µM) for 50 min, followed by a 10-min incubation with radioactive 2-deoxyglucose. Portions of the muscle extracts were analyzed by immunoblotting with the indicated antibodies (B-D, n = 5 per treatment/genotype; F-H, n = 4 per treatment/genotype); the remainder was used for glucose-uptake measurements (E, I, n = 4-5 per treatment/genotype). Data are representative of two independent experiments. Results are shown as mean ± SEM. Statistical significance was determined by two-way ANOVA with Tukey’s post hoc correction. \**P* < 0.05 (treatment effect within the same genotype), ^#^*P* < 0.05 (genotype effect within the same treatment).

## Supplementary Tables

**Supplementary Table 1.** List of primary antibodies.

| Antibody | Application<br>(IP/IB/IF) | Supplier | Reference |
| --- | --- | --- | --- |
| phospho-ACC<br>(Ser79) | IB | Cell Signaling | 3661 |
| ACC | IB | Cell Signaling | 3662 |
| phospho-Akt<br>(Ser473) | IB | Cell Signaling | 4060 |
| Akt | IB | Cell Signaling | 4691 |
| phospho-AMPKα<br>(Thr172) | IB | Cell Signaling | 2535 |
| AMPKα | IB | Cell Signaling | 2532 |
| AMPKα1 | IB | MilliporeSigma | 07-350 |
| AMPKα2 | IB | MilliporeSigma | 07-363 |
| AMPKβ1/β2 | IB | Cell Signaling | 4150 |
| AMPKγ1 | IP | Custom-made (YenZym)<br>The antibody was raised against the peptide<br>(CESSPALENEHFQETPESNNS, residues<br>E7–S26 of mouse AMPKγ1) | YZ7245 |
| AMPKγ1 | IB | AbCam | ab32508 |
| AMPKγ3 | IP/IB | Custom-made (YenZym)<br>The antibody was raised against a<br>combination of human and mouse AMPKγ3<br>peptide, human AMPKγ3 [residues S45–<br>S65: *CSSERIRGKRRAKALRWTRQKS],<br>mouse AMPKγ3 [residues S44–S64:<br>*CSSERTCAIRGVKASRWTRQEA] | YZ7243 |
| AMPKγ3 | IP | Custom-made (MRC PPU Reagents &<br>Services, University of Dundee)<br>The antibody was raised against a<br>combination of human and mouse AMPK $\gamma$ 3<br>peptide, human AMPK $\gamma$ 3 [residues S45–<br>S65: *CSSERIRGKRRAKALRWTRQKS], mouse AMPK $\gamma$ 3 [residues S44–S64:<br>*CSSERTCAIRGVKASRWTRQEA] | YZ7244 |
| phospho-STBD1<br>(Ser175) | IB | Custom-made (YenZym) | YZ7509<br>(PMID: 30772465) |
| STBD1 | IB | ProteinTech | 11842-1-AP |
| Vinculin | IB | Cell Signaling | 13901 |

IB; immunoblot, IP; immunoprecipitation

**Supplementary Table 2.** List of secondary antibodies.

| Name | Supplier | Reference |
| --- | --- | --- |
| Peroxidase AffiniPure Goat Anti-Rabbit IgG | Jackson ImmunoResearch | 111-035-144 |
| Anti-Mouse IgG HRP-linked | Cell Signaling | 7076 |
| Anti-Rabbit IgG HRP-linked | Cell Signaling | 7074 |

**Supplementary Table 3.** Quality control assessment of mouse snRNA-seq data.

| Sample ID | Number of nuclei | Median reads per nucleus | Median genes per nucleus | Total genes detected | Median UMIs per nucleus | Mitochondrial genes (%) |
| --- | --- | --- | --- | --- | --- | --- |
| 1 | 2398 | 2351 | 1317 | 1321 | 2351 | 0 |
| 19 | 3823 | 2015 | 1269 | 1506 | 2015 | 0 |
| 2 | 1047 | 2070 | 1130 | 632 | 2070 | 0 |
| 20 | 1339 | 973 | 600 | 702 | 973 | 0 |
| 21 | 4206 | 3128 | 1536 | 2207 | 3128 | 0 |
| 3 | 1667 | 1821 | 1231 | 967 | 1821 | 0 |

**Supplementary Table 4.** Quality control assessment of human snRNA-seq data.

| Sample ID | Number of nuclei | Median reads per nucleus | Median genes per nucleus | Total genes detected | Median UMIs per nucleus | Mitochondrial genes (%) |
| --- | --- | --- | --- | --- | --- | --- |
| 11-0 | 113 | 2290 | 1348 | 110 | 2290 | 0 |
| 11-8 con | 2744 | 2594 | 1629.5 | 1548 | 2594 | 0 |
| 13-0 | 854 | 2269 | 1510.5 | 634 | 2269 | 0 |
| 13-8 con | 1216 | 1839 | 1278.5 | 840 | 1839 | 0 |
| 14-0 | 3094 | 1922 | 1032 | 849 | 1922 | 0 |
| 14-8 con | 1952 | 2941 | 1702.5 | 1165 | 2941 | 0 |
| 26-0 | 2062 | 2858 | 1855 | 1240 | 2858 | 0 |
| 26-8 con | 2655 | 2428 | 1537 | 1499 | 2428 | 0 |
| 28-8 con | 1994 | 1650 | 1078 | 1042 | 1650 | 0 |
| 31-0 | 1877 | 3218 | 1931 | 1213 | 3218 | 0 |
| 31-8 con | 2031 | 3416 | 1899 | 1348 | 3416 | 0 |
| 32-0 | 3530 | 3375 | 1874 | 1661 | 3375 | 0 |
| 32-8 con | 3572 | 3258 | 1829 | 1829 | 3258 | 0 |

## Notes

### Competing Interest Statement

The authors have declared no competing interest.

